# Ex vivo Infrared Nerve Stimulation on the Rat Sciatic Nerve: Challenges and Pitfalls

**DOI:** 10.64898/2026.03.06.709613

**Authors:** Carlos Izquierdo Geiser, Christian Münkel, Paul Schlett, Patrick Campbell, Gergana D. Borisova, Celine Wegner, Karin Somerlik-Fuchs, Ulrich G. Hofmann

## Abstract

Infrared nerve stimulation (INS) offers millisecond-scale, electrical artifact-free activation of peripheral nerves, yet rat ex vivo studies remain scarce. We present an INS setup utilizing a lens system for free-beam focus for rat sciatic nerves partially submerged in a nerve bath, making re-wetting of the tissue in existing ex vivo INS setups redundant. Excised sciatic nerve preparations were illuminated at 1470 nm, delivering pulse trains from 100 – 2000 μs at 5 Hz with radiant exposures up to 25.7 J/cm^2^. Compound action potentials (CAPs) were recorded with tungsten hook electrodes. Nerve activity could be recorded up to 200 minutes after excision, though persistent nerve excitability remained comparable to existing ex vivo studies as a yet to be optimized parameter. Peak CAP amplitudes ranged from 3.9 - 36.9 µV with latencies averaged at 3.4 ms. Activation thresholds spanned 1.75 – 13.05 J/cm^2^.

Additionally, potential pitfalls for INS setups and ways to resolve them were investigated, with two artifact types highly relevant to ex vivo INS being identified: A photo-thermal expansion artifact and a photovoltaic artifact with thermo-capacitive coupling.

The observations confirm that CAPs can be evoked via INS in an ex vivo preparation of the rat sciatic nerve. The presented platform supports 3R principles and offers a robust basis for introduction of pharmacological investigations with INS.

## Introduction

Neuromodulation is an established clinical approach used in the peripheral nerves in various applications such as neuronal conduction velocity testing, motor stimulation in patients with diaphragmal palsies, hemi-, and paraplegia, and sensory nerve stimulation for neuropathic pain control (Slavin 2011, Takeda et al. 2017, Ganji-Angirekula & Javed 2024). While functional electrical stimulation (Bulley et al. 2021) and transcranial magnetic stimulation (Deng et al. 2025, Zhi et al. 2025) are widely and actively used in clinical practice, multiple other promising approaches have been investigated, such as stimulation by ultrasound (Feng et al 2019) or illumination at various light wavelengths (Martings et al. 2025, Zhi et al. 2025). These alternative modalities aim to expand the repertoire of clinical tools available to clinicians and researchers, each bringing its advantages and challenges.

Infrared nerve stimulation (INS) is one such promising alternative. Initially described in 2005, the approach utilizes irradiation of genetically unmodified nerves directly with laser pulses in short time frames to elicit nerve activity (Wells et al. 2005, Wells et al. 2005b). Its characteristic advantages lie in the high temporal and spatial resolution of stimulation, preventing unwanted recruitment of other neural tissue in the vicinity. This level of control is difficult to achieve with conventional electrical or magnetic methods, which tend to recruit further excitable structures within a broad volume. Furthermore, INS is inert to many other imaging applications such as magnetic resonance imaging and especially other electrophysiology techniques since the nerve activation is free of electrical artifacts, allowing for an unaltered observation of the elicited compound action potentials (CAP). (Wells et al. 2005, Chernov & Roe 2014, Song et al. 2024)

The mechanism of INS is yet to be determined in its entirety (Chernov & Roe 2014, Song et al. 2024), and damage thresholds as well as the reliability of stimulation remain active areas of research. Another characteristic of INS is that each nerve is reported to have individually located regions of interest (ROIs) that are highly susceptible to stimulation by laser irradiation, while other areas of the same nerve do not elicit any reaction to the same stimulus (Duke et al. 2012, Peterson & Tyler 2015).

Typically, INS studies are reported to have been carried out in vivo. Alternative methods such as in vitro neural organoids are found to be challenging due to their high heterogeneity and incomplete replication of organ function (Su et al. 2025). Yet, with animal welfare taken seriously, the 3R principles (replacement, reduction, refinement) gain increasing importance and a demand for viable alternative experimental methods and setups to reduce stress and suffering of animal models is tangible (Neziri et al. 2024, Zhang et al. 2024). Utilising ex vivo models with excised tissues would allow for increased throughput of experiments, reduced compliance challenges regarding ethical issues, and maintains the physiological and functional cohesiveness of the nerve fibre (Lossi & Merighi 2018). By conducting experiments on isolated nerves, researchers can apply precise, repeatable stimuli while avoiding systemic effects that confound interpretation in whole animals. An ex vivo approach using excised nerve tissue, such as the sciatic nerve of rats, would clearly permit multiple experiments on different tissue segments, and could thereby reduce the overall animal burden in line with the 3R’s.

Ex vivo studies are particularly advantageous because they allow the introduction of compounds modulating nerve function otherwise affected by anesthetics mandated during in vivo experiments (Rapeaux et al. 2022, Berkowitz et al. 2025). Furthermore, in contrast to full animal settings utilized pharmaceuticals can be reliably washed out (Rapeaux et al. 2022). This precise chemical control might be particularly valuable when testing time sensitive agents targeting neural tissue.

Despite these advantages, ex vivo work is rarely employed for INS and is in rodents largely limited to electrical stimulation (Sun et al. 2020, Rapeaux et al. 2022, Berkowitz 2025). To our best knowledge, only two rat ex vivo INS setups were established, which were utilized for a wavelength INS study (Katz et al. 2009), and for conduction velocity and temperature measurements (Cury et al. 2021, Perre et al. 2022, Perre 2025, Perre 2025b). As the former setup is based on gravity perfusion and the latter setup does require repeated wetting of the nerve, a gap remains for the active and seamless introduction of pharmacological compounds that alter nerve functionality in INS setups.

The study at hand aims to validate an ex vivo setup that largely follows mentioned established electrical stimulation approaches. By adapting this platform to INS, we aim to provide an ex vivo method that promotes the 3R principles. In doing so, we seek to bridge the current gap between in vivo efficacy studies and the need for more refined, animal sparing experimental designs.

## Methods

### A. Setup and Tissue Preparation

INS stimulation was performed with a commercial fibre-coupled infrared diode laser system (1470 nm, 30 W, 105 μm, NA 0.22, Aerodiode CCM-I 1470LD-5-4-2, and 3 m fibre 105 μm, NA 0.22, AeroDIODE, Bègles, France) projecting through a custom designed, focusing lens system (Fig. 1. A and Supplemental Material 2). Position and distance between nerve and lens system across x-y-z-axes can be adjusted by a 3-axes micromanipulator (Mini 25, Luigs & Neumann GmbH, Ratingen, Germany) with ±0.1 μm accuracy. Interfacing and control of the setup is carried out by custom python programs running on a desktop PC. Arduino UNO microcontrollers are utilised for synchronisation and triggering. Neural activity of the sciatic nerve is detected with 100 μm diameter tungsten wire hook electrodes (curvature radius 1.5 mm) with help of a digitizing headstage (RHD 32-Channel Recording Headstage, Intan Technologies, Los Angeles, USA) and the Open Ephys DAQ acquisition board (Open Ephys Inc., Atlanta, USA, Siegle et al. 2017). The remaining setup follows in principle the one described for electrical stimulation by Rapeaux et al. 2022.

**Fig. 1:**
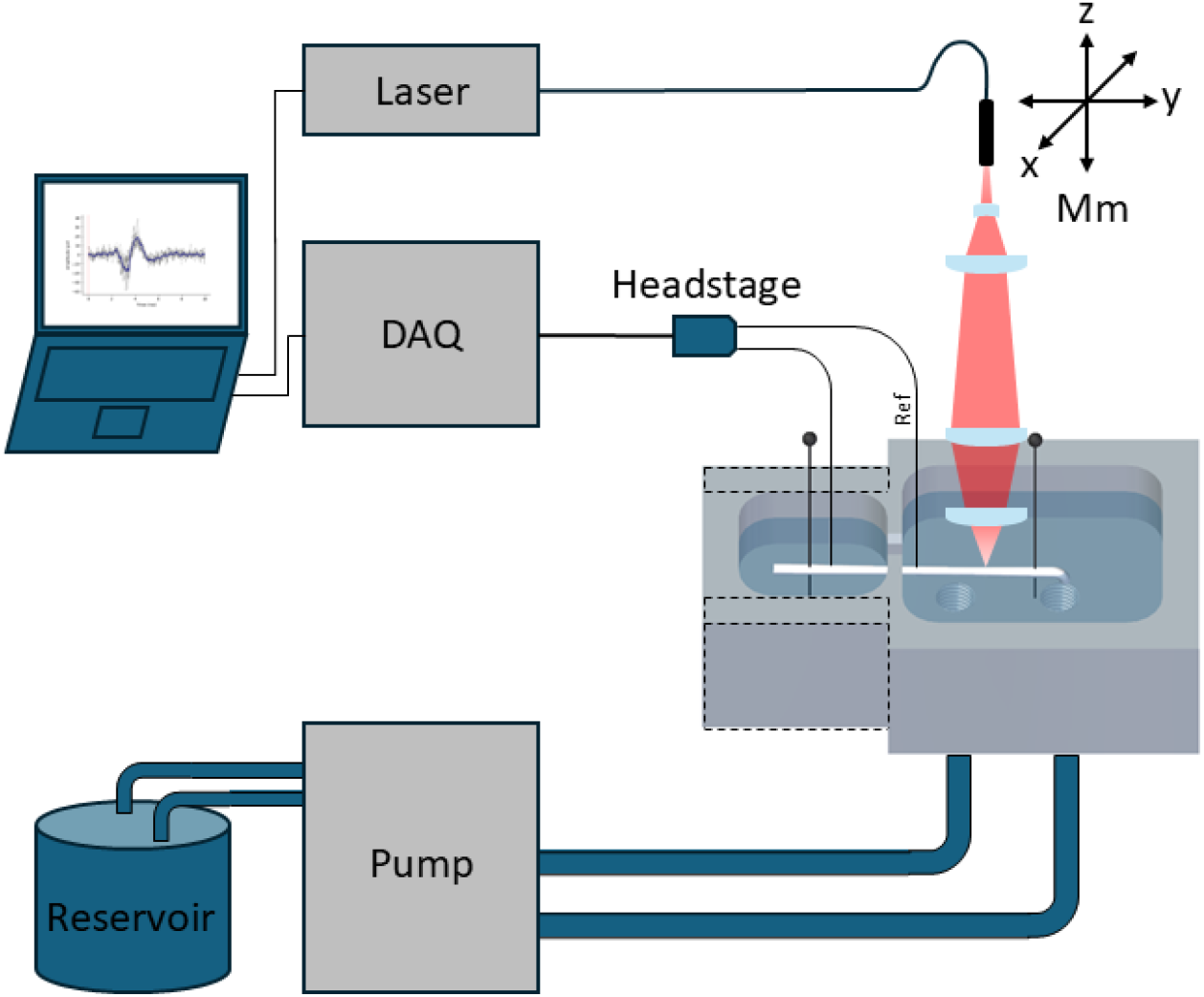
Experimental Setup. The free laser beam exiting the primary glass fiber is focused through a lens system positioned in all three axes with a PC controlled micromanipulator (Mm). An excised rat sciatic nerve is mounted in a nerve bath, with detachable side walls for better mounting accessibility (dashed lines), permanently supplied with buffer solution via a pump at 4 ml/min. Data is collected by hook electrodes over a headstage connected to the Open Ephys DAQ.

Tissue sourcing was performed with approval from the local Animal Welfare Committee at the Regierungspräsidium Freiburg in accordance with the guidelines of the European Union Directive 2010/63/EU (permit X/21 02R). Adult male Sprague-Dawley rats (N = 9, 300 - 500 g, 8 – 12 weeks old, Charles River WIGA, Sulzfeld, Germany) were housed in groups with food and water available ad libitum. Rats were anaesthetized with Isoflurane (4-5 % in O2, Forene, Abbot GmbH & Co. KG, Wiesbaden, Germany) and subsequently injected i.p. with ketamine (100 mg/kg) and xylazine (8 mg/kg) until being reflex free. Afterwards the rats were sacrificed by immediate decapitation allowing use of multiple organs for other experiments. The skin was shaved over the thighs, and the sciatic nerve was bilaterally exposed by incision of the skin and the muscular fascia on both sides. This was followed by a blunt dissection between the gluteus superficialis and biceps femoris, which was held open by a surgical retractor and moistened frequently with saline solution until excision. The sciatic nerves of 2 - 3 cm length proximal to the bifurcation were subsequently extracted and placed into chilled oxygenated mKHB (see below) where they rested until experimentation.

A physiological modified Krebs-Henseleit buffer (mKHB) solution (200 ml mKHB, 2.1 g sodium bicarbonate, 0.99 g dextrose, 1.8l deionised water, carbogenisation (5% CO_2_, 95% O_2_) for at least 30 minutes at 40 °C) is used to maintain the nerve for extended viability. Heating the reservoir to 40 °C results in a temperature of 37° C after pumping (PPS2, Multichannel Systems MCS GmbH, Reutlingen, Germany) it into the nerve bath chamber. A commercial temperature probe (TSP01, Thorlabs, Newton, USA) is used to control the buffer reservoir after initial calibration of the temperature difference between reservoir and nerve bath. In- and outflow sources have been modified from the original protocol to achieve a more consistent and uniform distribution of the buffer solution. The setup is schematically depicted in Fig. 1. An excised sciatic nerve is mounted in a 3D printed biocompatible nerve bath. It holds the intact nerve with two pins in two sub-chambers where one is used for recording and the other as stimulation site, respectively. After mounting the nerve with pins into both chambers, the connecting orifice is sealed with Vaseline (Balea, Aachen, Germany). In addition, the recording chamber is filled with mineral oil (Carl Roth, Karlsruhe, Germany) to counteract volume conduction artifacts from one chamber to the other and thus ensures proper recording of nerve activity. The mKHB solution is recirculated between reservoir and stimulation chamber with a rate of 4 ml/min.

The sidewalls (Fig. 1, dashed line) were designed to be detachable, giving easier access to the nerve in the recording chamber and thus reducing mechanical stress while attaching the electrodes. Before infrared illumination nerve viability was tested by electrical stimulation (BCS-probe and ISIS Neurostimulator, inomed Medizintechnik GmbH, Emmendingen, Germany) via a micro manipulator positioned bi-polar probe on the nerve.

### B. Laser Beam Characterization

A lens system was designed and implemented to enable contactless irradiation of the sciatic nerve with a focused, symmetric, free laser beam instead of the commonly used stimulation from the working end of an unfocused fibre. To enable consistency and comparability with existing literature radiant exposure H_e_ [J/m^2^] (often reported in J/cm^2^) is used for describing the energy deposition onto the target tissue. For unambiguous description of the parameters thus the pulse energy E_p_ [J], and the diameter of the focal spot size d [m] is needed. Due to the curved surface and anatomical variations of each nerve, an exact spot size is most difficult to determine. Therefore, our free laser beam is characterized prior to experiments via a rotating-slit infrared beam profiler (BP209-IR/M, Thorlabs, Newton, NJ, US). The beam waist is determined to be d = 2w_0_ = 352 μm, which results in a spot size of 9.73·10^-4^ cm^2^. Pulsed laser output power, measured by a pyroelectric pulse energy sensor (ES145C, Thorlabs, Newton, NJ, US), was operated at the maximum available laser output power. The constant optical power output enables linear adjustment of deposited pulse energy through adjusting illumination time (pulse width) up to 2 ms resulting in a maximum radiant exposure of 25.71 J/cm^2^.

The lens system’s caustic is designed such that the beam achieves depth of focus of d_w_ = 1.5 mm within which the beam radius remains below 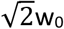 at a focal distance of d_f_ = 2.1 cm from lens to nerve surface (Fig. 3 A). The accurate distance to the nerve is ensured by a 3D printed soft tip stylus attached to the front of the lens system.

### C. INS Protocol

For each stimulation a standardized train of pulse sequences is applied. Pulse sequences consist of 10 pulses at a set pulse energy E_p_, and a pulse width τ_p_ at a frequency of f_p_ = 5 Hz with 2 seconds between pulse sequences to counteract temperature buildup. Pulse trains consisted of 20 pulse sequences at steadily increased equidistant pulse widths (Fig. 3 D) by which the radiant exposure H_p_ is increased over the course of the pulse train. The sciatic nerve is stimulated between the pelvic exit point and the bifurcation of the medial section. Laser position is varied between pulse trains over the course of the experiment within the constraints of the described nerve area. Locations are chosen at random without testing or consideration of whether the area is a presumably easy to stimulate ROI or non-ROI. A distance of ≥1 mm was maintained between each stimulation spots, as measured by the micromanipulator.

### D. Data analysis

All pulse sequence parameters are noted for the experiments, while electrophysiological data is recorded at a sampling rate of f_s_=30 kHz continuously via the Open Ephys GUI. Control measurements are performed before experimentation without any stimulation to acquire baseline noise data V_N_. Data analysis is performed using a customized Python program. Incident laser pulses are marked directly in the data via TTL-signals to identify CAP in the following 20 ms window. If the mean signal of one pulse sequence (10 single pulses) of tentative CAPs exceed the positive or negative threshold value of two times the root-mean-square value (RMS, threshold = 2·V_N_) it is marked as successful elicited nerve activity. A 50 Hz notch filter against typical noise from power lines is applied. Negative controls with occluded beam path are carried out to ensure the absence of crosstalk artifacts and proper function of the system.

## Results

### Successful CAPs are produced by short laser illumination ex vivo

Although a total of 9 Sprague Dawley rats were used in this study only 14 sciatic nerves were usable for the experiments. The remaining nerves were discarded due to either unsuccessful surgery, or in one case each due to unresponsiveness to electrical stimulation, and tissue damage in the mounting process of the nerve, respectively. pH values of the buffer solution were stable across the whole duration of experiments at 7.37 ± 0.11. The sciatic nerves were mounted so that all but the top side was submerged in the buffer solution to ensure adequate supply of nutrition to the nerve but not interfere with the light beam. The reference and recording electrode were separated by 11.5 mm. As a correct and good connection to the nerve tissue took precedence, slight differences in the desired distance between the electrodes was expected and observed. With the spot diameter size of d = 2w_0_ = 352 µm and a nerve diameter of approximately 1 – 1.5 mm sufficient area for proper position and scanning was given. Infrared laser pulse trains were able to elicit in 8 nerves out of 5 rats a measurable CAP (Fig. 2, for pulse groups average see Supplemental Material 1). Successful CAP responses were observed while scanning a single nerve up to 35 minutes post excision. As no parallel experimentation on two excised nerves was possible, the second nerve rested significantly longer in carbogenized medium than the first one from each particular rat. This led to a maximum post excision delay of 200 minutes for a still successful stimulation (as defined by detecting CAP). Baseline RMS noise over all measurements was observed to be 4.58 ±3.36 µV. Due to the overall small amplitude of the evoked CAPs all further analyses are performed on averaged signals for each pulse sequence to decrease random noise influences.

**Fig. 2:**
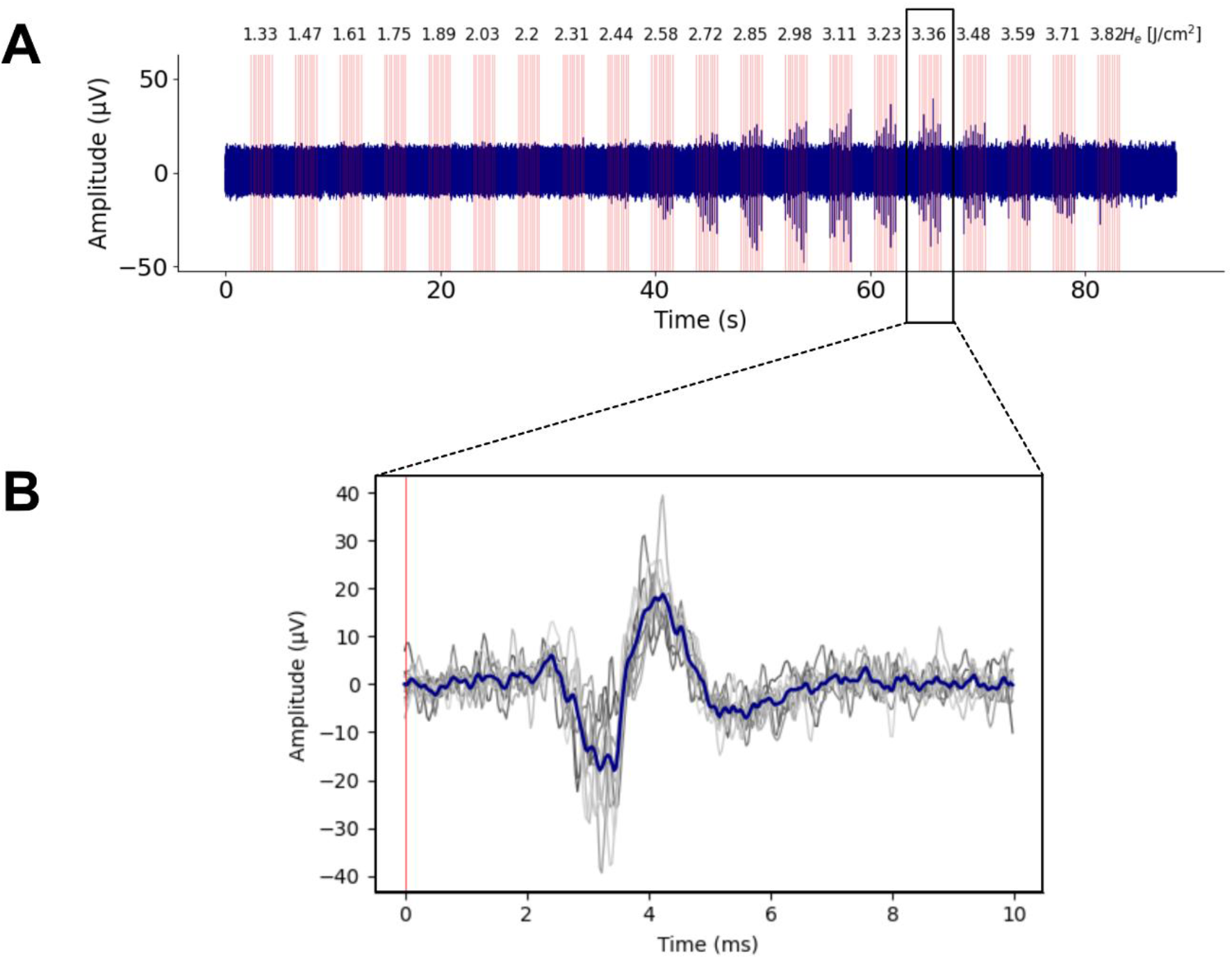
A) Representative results for one full experiment run with single pulses marked by red lines. Radiant exposure H_e_ is successively increased with each pulse sequence. B) Observed representative CAP at 3.36 J/cm^2^ of 10 pulses (grey) and mean (blue). Red line marks end of incident laser pulse.

A low signal to noise ratio of 8.91 dB (average at maximum average amplitude per pulse train) was observed, making the accurate CAP onset at lower amplitudes hard to distinguish on the noisy background. This and the geometric nerve variability resulted in an average latency for detecting CAP responses of 3.37 ± 0.63 ms.

Maximum CAP amplitudes ranged from 3.92 – 36.91 µV (single results seen in Tab. 1) with typical signal forms depicted in Fig. 2 B. The CAP amplitude appeared to monotonously increase to a maximum along successively increasing radiant exposure. Exceeding a certain radiant exposure then led to a steady decrease of the measured voltage modulation until no signal was observed anymore (Fig. 2 A). The initial increase and subsequent decrease showed a large variability with the smallest values seen over a range of 3.23 to 3.82 J/cm^2^, while the largest was observed over a range of 10.01 – 25.71 J/cm^2^. Furthermore, in the latter case the amplitude was still visible at the maximum pulse width of 2 ms suggesting a wider range beyond the scope of the stimulation parameters used.

**Fig. 3:**
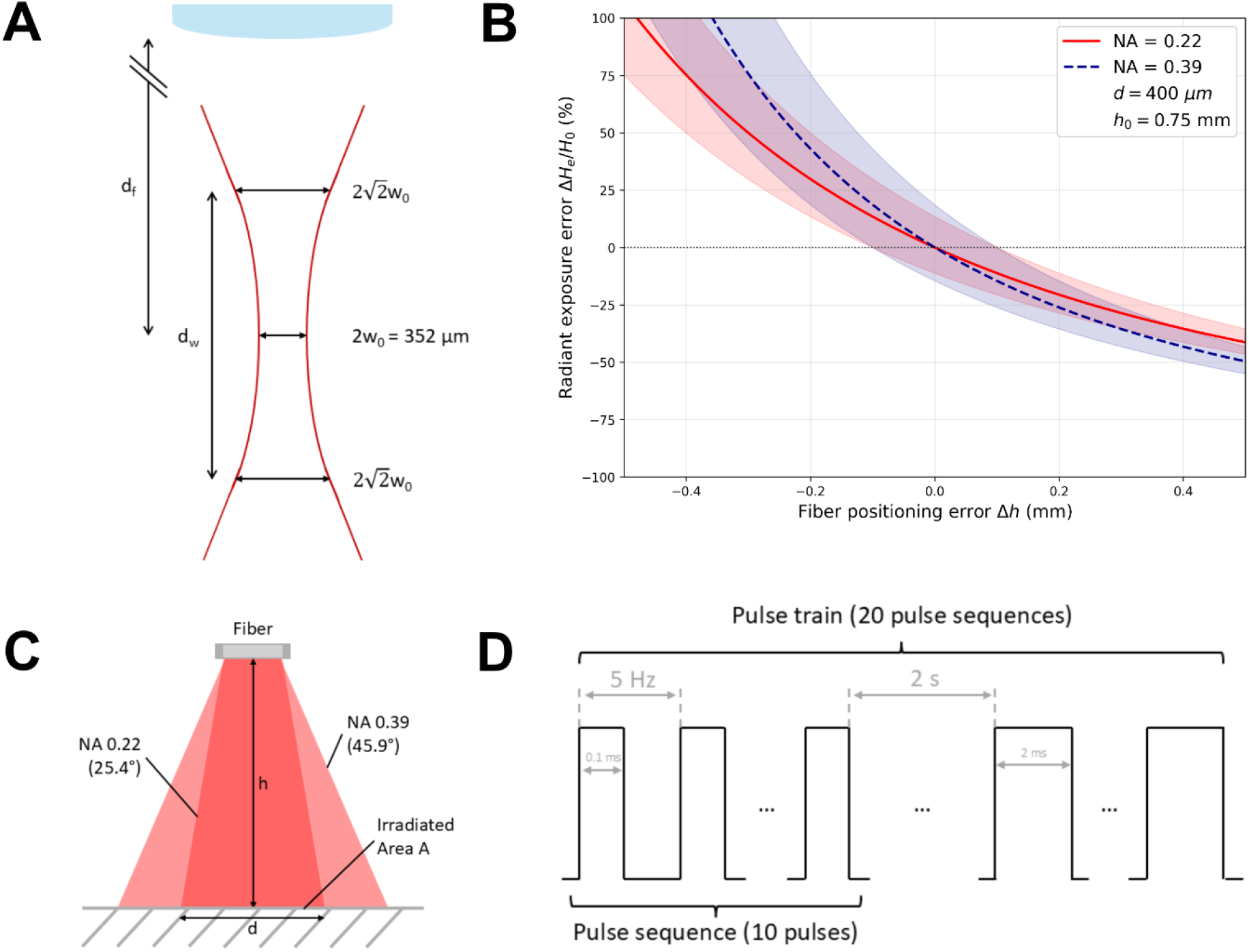
A) With our simple lens arrangement the free beam forms a projection caustic with a given beam waist 2w_0_ = 352 μm varying less dramatically over its useful working depth of focus d_w_ at a distance d_f_ from lens to focus. B) The illuminated area from a bare glass fiber end is crucially dependent on the precise distance towards the neural target, due to the opening angle of the illumination cone given by the fiber’s numerical aperture. Systematic fibre positioning error due to fibre misalignment is shown for the example of a 400 µm fibre with NA = 0.22 (red) or NA = 0.39 (blue) at an assumed distance of 0.75 mm above target. CI depicts positioning accuracy of ±100 µm showing the impact of minor misalignments due to unforeseen positioning mistakes or movement. C) Schematic visualization of exit cones of optical fibers and the spot size d depending crucially on the height h from the surface and cone angle defined by the numerical aperture NA. D) Schematic overview of applied pulse trains each consisting of 20 sequences with a pause of 2 ms between each. Each pulse sequence consists of 10 pulses at 5 Hz ranging from 0.1 ms to 2 ms. Pulse width is increased by 0.1 ms for each new pulse sequence.

### Free beam illumination approach to minimize geometrical projection influences

Commonly, illumination for INS experiments are performed simply with an optical fibre, where the fibre is placed in an optimistically defined position towards a nerve. However, due to the divergence of light exiting from bare fibre ends (typically with a numerical aperture of NA = 0.22 or NA = 0.39, schematically visualized in Fig. 3 C a minor shift in actual distance in relation to the target results in a relatively large increase in beam cone diameter. Considering that the radiant exposure strongly depends on the illuminated area, factoring into the beam radius with the power of two, such a change in radius may have a large impact on the deposited energy into the nerve. Furthermore, the influence for different height aberrations in Fig. 3 B and assuming a minute discrepancy of 100 µm in positioning of an exemplary 400 µm fibre with an NA = 0.22 or NA = 0.39, respectively, results in a 13.5 % (NA = 0.22) or 18.6 % (NA = 0.39) error in radiant exposure. Since the higher NA results in a wider emission cone a larger impact on this error is not unexpected. Larger distances between fiber end and target may therefore result in a lower applied radiant exposure on the nerve, while an accidental shorter distance may lead to a potentially damaging over-exposure. Under-exposure may then experimentally be compensated by stronger laser pulses but will then decrease the spatial resolution and introduce unwanted damage. All of these simple geometrical considerations are further confounded by the nerve diameter being potentially smaller than the illumination spot, thus “wasting energy” or even illuminating adjacent tissue facilitating potentially damage to those regions.

We thus decided to use a free-beam laser illumination which is, due to its 1.5 mm long caustic, resistant to small changes in the geometrical relation between beam and nerve.

### Photo-thermal expansion may be erroneously interpreted as CAP

Since we set out to thoroughly characterize INS, a careful evaluation of apparent stimulation successes caused us to widen our set of potential confounding influences. Fig. 4 A displays a seeming prime example for a sought-after event: The hook electrodes pick up tentative CAPs after illuminating an ex vivo sciatic nerve with 10 pulses, each with 100 µs duration at 1.81 J/cm^2^.

**Fig. 4:**
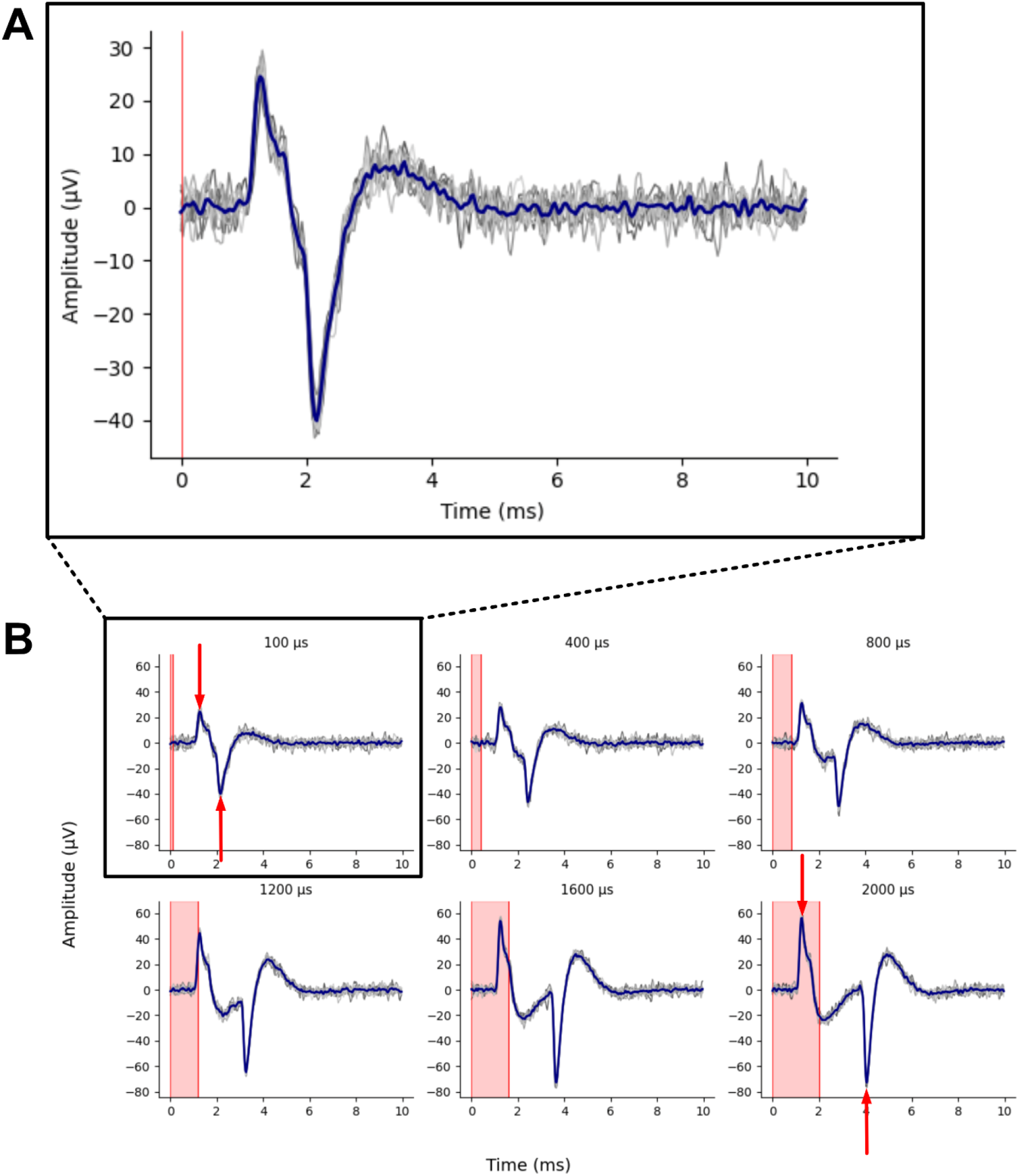
A) 10 ms windowed measurements depicting mechanical artifacts (grey – single measurement, blue – mean of pulse sequence) induced by the incident laser beams in pulse trains (10 pulses, 5 Hz, 100 – 2000 µs pulse width, blue line marks mean), which have a striking similarity with CAP signals at small pulse widths. B) At successively increased pulse widths it becomes apparent that the potential CAP is an artifact and the whole signal consists of two superimposed identical but inverse parts that occur at the beginning and end of the laser beam. The artifact is furthermore observable while the laser pulse is still irradiating the nerve.

In some cases with relatively short (∼2 cm) nerve explants, the nerve was positioned as usual between the fixating pins and appeared to feature a recognizable tentative CAP upon laser illumination (Fig. 4 A). CAPs after electrical stimulation (positive control) ensured each time the viability of the nerve also in these specific cases. However, its artifactitious character was initially discovered when finally and reliably severing the nerve conduction by nerve-crush with a tweezer: this signal was still visible upon illumination. This artifact, encountered in our experiments at 100 µs single pulse width, follows the commonly taught form of a nerve excitation (Crone and Krarup 2013) which on its own could possibly lead to the wrongful assumption of a physiological nerve response to INS. Its amplitude of the tentative CAP is indeed increasing with applied pulse width and hence deposited energy. When observing the development of the signal over increasing pulse length (Fig. 4 B), however, it becomes apparent that the biphasic signal consists of two superimposed, mirrored spikes. With longer illumination pulse length, the two mirror spikes gain temporal separation to each other (red arrows, Fig. 4 B). The apparent artifact becomes more prominent with longer pulses and appears eventually while the illumination is still active. This is another sign of a non-physiological artifact, as proper INS produces actual CAP signals with a physiological delay following the end of incident laser pulses [Wells et al. 2007].

Additionally, this effect was also observed if the laser beam was slightly misaligned and thus partially missed the nerve. In this case, similar artifact signal forms were observed though with even higher magnitude. The incident beam caused visible waves forming in the medium, which were accompanied by an audible sound.

This artifact could for the rest of the study be avoided, as mechanical tension on the nerve could be reduced by positioning pins slightly closer to each other. Furthermore, longer nerves could always be positioned with minimal tensions not expressing this artifact.

### Light-electrode interaction may be erroneously interpreted as CAP

Once sensitized to the potential pitfalls of the light-medium interaction, we searched for other ways to introduce erroneous CAP events. One way of reproducibly acquiring such artifacts was by illuminating the close proximity (<2 mm, Fig. 5 B) of the recording electrodes. Changing our usual tungsten hook electrodes to platinum or illuminating it without the sciatic nerve made qualitatively no difference: We reliably saw transient signals in the range of several 10 µV, whether or not a nerve was mounted on the hook.

**Fig. 5:**
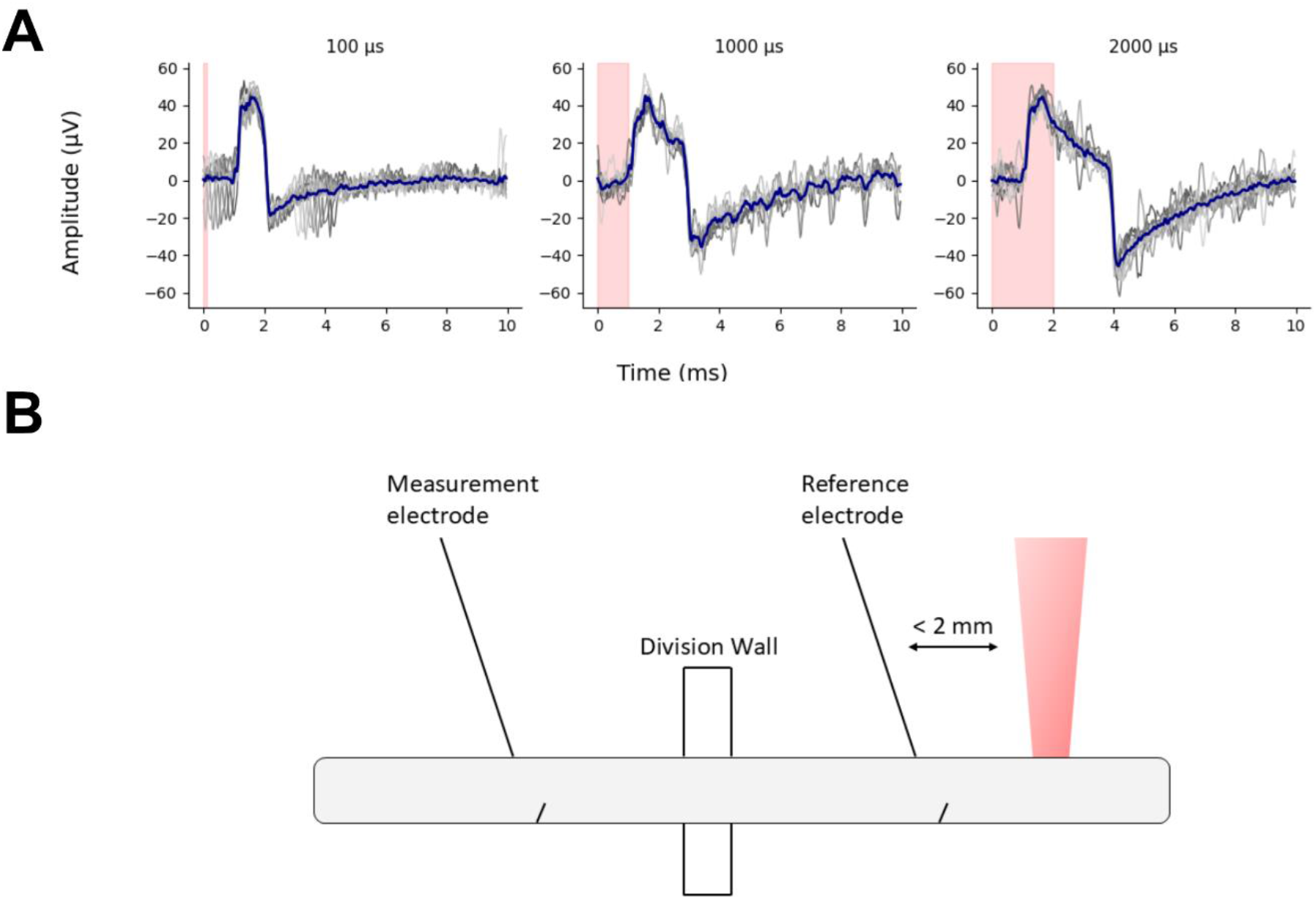
A) Observed artifact caused by illumination in close proximity to recording electrodes (Tungsten 100 μm). Red area shows active incident laser pulse. Rising flank of the artifact is measured 1.1 ms after the initial laser pulse, while the falling flank occurs 2.1 ms after cessation of the laser pulse. The following signal rises afterwards slowly over multiple milliseconds back to the baseline of 0 μV. B) Geometry of the setup, nerve, and incident laser beam resulting in the observation of the light-induced artifact.

This artifact (Fig. 5 A) is characterized by sharp flanks and a stable maximal amplitude irrespective of the induced laser energy. All such observed artifacts amounted to the same amplitude of approximately 40 µV. From the initial fast rise, 1.1 ms after the first laser pulse, the signal decays monotonously until a fast drop of approximately -60 µV trails the last laser pulse by 2.1 ms. All in all, peak-to-peak amplitudes ranged between 60 µV and 90 µV. Given some “recuperation time”, the signal returns after a few milliseconds back to the base level of 0 µV with longer times being needed at higher pulse width due to larger negative amplitudes. The leading parameter influencing the peak-to-peak amplitude was the time span the laser illumination was active and with this its deposited energy.

## Discussion

This study was conducted to validate our free-beam setup for INS in at 1470 nm wavelength. The illuminated nerve is resting in a buffer solution, and the elicited CAPs are well detectable, even more so with averaging over 10 individual pulses of one pulse sequence (Fig. 2 B, blue). Amplitudes of individual traces within each stimulating sequence (Fig. 2 B, grey) showed variability due to the influence of noise on these very low amplitude signals. Because the signal is so small, even minute fluctuations in the background noise have a relatively strong impact producing noticeable distortions in the measured CAP. The setup demonstrates similar success rates of observed CAPs as the existing literature (Cury et al. 2021). Our setup differs from other INS ex vivo setups which rely on continuously wetting the nerve with a syringe pump (Perre et al. 2022) or just drop-wetting it in certain time intervals (Cury et al. 2021). In order to maintain conditions as constant as possible over the span of each experiment, we used a nerve bath allowing the excised neural tissue to stay mostly submerged in a stable environment. It fosters general hydration and thus avoids strong variations in wetting layer thickness on top of the nerve, potentially absorbing light with a high variability before it may interact with the actual nerve tissue (Wells et al. 2007a). Still, despite our best efforts and comparable CAP values to literature throughout, biological variability remained high, as one rat elicited CAPs only at substantially elevated energy depositions (Tab. 1).

**Tab. 1:**
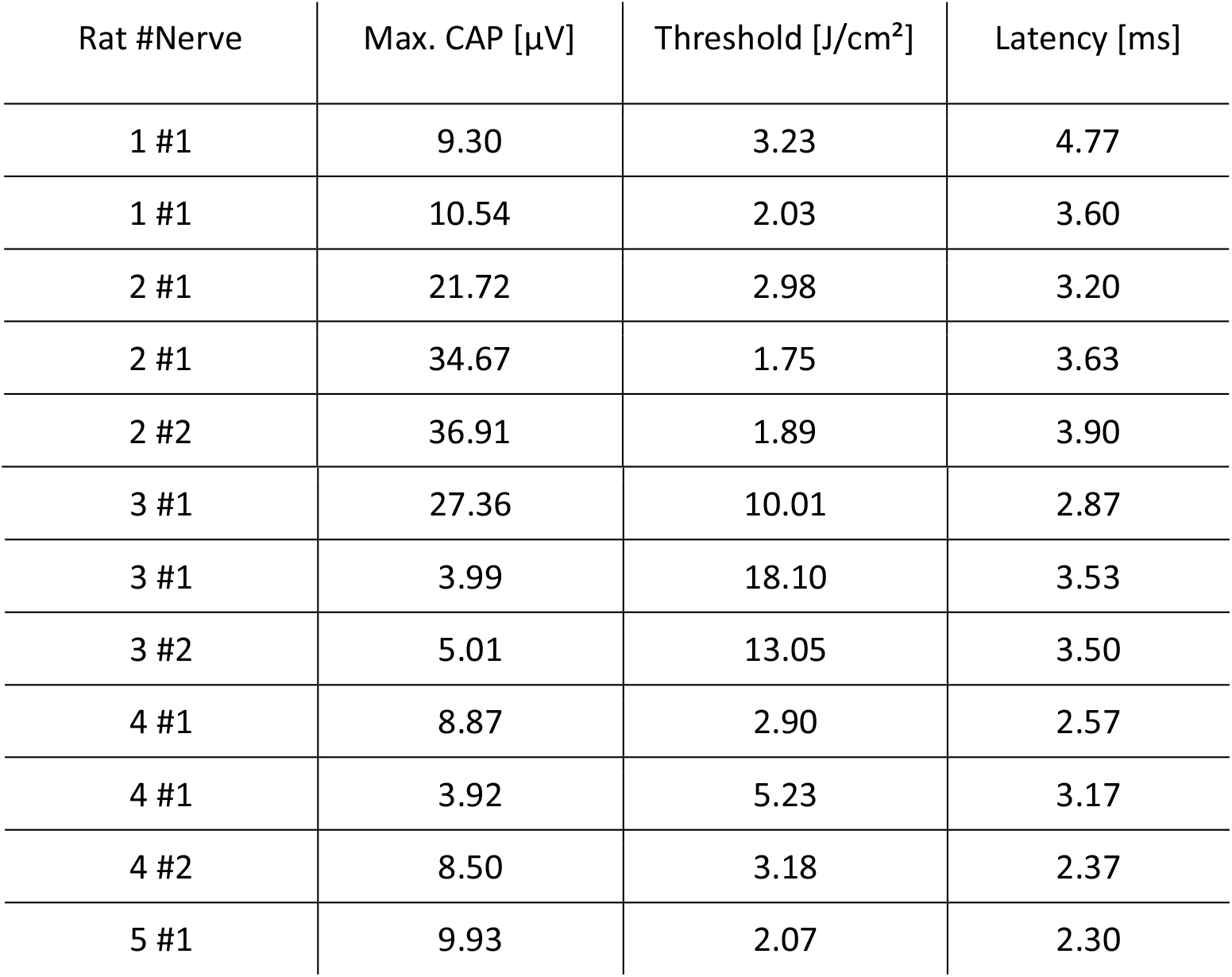
Overview of all measured experiment values per rat and nerve.

| Rat #Nerve | Max. CAP [$\mu\text{V}$] | Threshold [$\text{J}/\text{cm}^2$] | Latency [ms] |
| --- | --- | --- | --- |
| 1 #1 | 9.30 | 3.23 | 4.77 |
| 1 #1 | 10.54 | 2.03 | 3.60 |
| 2 #1 | 21.72 | 2.98 | 3.20 |
| 2 #1 | 34.67 | 1.75 | 3.63 |
| 2 #2 | 36.91 | 1.89 | 3.90 |
| 3 #1 | 27.36 | 10.01 | 2.87 |
| 3 #1 | 3.99 | 18.10 | 3.53 |
| 3 #2 | 5.01 | 13.05 | 3.50 |
| 4 #1 | 8.87 | 2.90 | 2.57 |
| 4 #1 | 3.92 | 5.23 | 3.17 |
| 4 #2 | 8.50 | 3.18 | 2.37 |
| 5 #1 | 9.93 | 2.07 | 2.30 |

Additionally, the activation thresholds encountered in our study are with a minimum of 1.75 J/cm^2^ significantly higher as compared to in vivo thresholds of 0.32 - 0.4 J/cm^2^ (Wells et al. 2005, Throckmorton et al. 2021, Wells et al. 2007) in rat sciatic nerves. Our average thresholds are on par with INS ex vivo literature which reports values of 3.16 ± 0.68 J/cm^2^ (Cury et al. 2021), though further research is necessary to shed light on the differences in mode of operation and underlying mechanisms of ex vivo and in vivo INS that would explain this consistent gap of approximately one order of magnitude for the activation threshold.

Our ex vivo study did not succeed to elucidate the actual mode of operation of INS, reportedly based on the workings of several temperature sensitive ion channel types (Suh et al. 2009, Albert et al. 2012). In order to do so, we would have to introduce specific ion channel blocker, chemically modulating CAP production. Unfortunately, appropriate comparative testing could not be conducted as the longevity in terms of re-excitability of the nerve was not sufficient, which, considering the literature, seems to be a ubiquitous and worthwhile challenge with ex vivo INS. Only a small subset of nerves elicited a second successful set of of CAPs irrespective of whether the location was established earlier or fresh on the same nerve, both directly after a previous stimulation or with 10 minutes pause to allow for some recuperation. This comes in stark contrast to our electrical ex vivo stimulation efforts, as in the same setup the excised nerve was successfully and repeatedly electrically stimulated for up to 60 minutes (data not shown).

It has to be noted that one of the advantages of the proposed ex vivo setup is its potential to comply with 3R standards by using the sciatic nerves from other acute experiments. In addition, it allows to investigate artifacts potentially obscured by anesthetized animal’s movement and -in principle-allows neuropharmacological investigations.

Noteworthy, while characterizing our minimalistic INS experiment, we came along a set of erroneous CAP-looking signals turning out to be artifacts, worth describing for widespread consideration in future studies.

The first artifact is introduced in Fig. 4 and can be produced by causing mechanical changes in the nerve or the nerve bath liquid. As we keep the running laser energy at 100% but modulate the radiant exposure by adjusting the pulse width of each of the 10 incident pulses between 100 – 2000 µs we successfully decomposed the apparent CAP in relation to time and energy. With lower pulse width, a biphasic pulse of approximately 3-4 ms is displayed, resulting in signals resembling a CAP (Fig. 4 A). However, the artifact sets itself apart from CAPs as it is already visible while the tissue is still being irradiated by the laser pulse. The difference to nerve activity becomes clear while increasing radiant exposure by longer active laser times, as the artifact’s signal decomposes and components shift apart (Fig. 4 B). The signal shape does change its amplitude with increased pulse widths, similar to physiological INS CAPs. However, we speculate that the increase in amplitude is not a physiological CAP but rather caused by stronger photo-thermal induced mechanical influences. This artifact may occur at multiple stimulation loci irrespective of their distance to the electrodes, distinguishing it from the photo-electric artifact on electrodes, described further down. As mentioned, the main evidence for a mechanical artifact are the audible and visible waves in the nerve bath upon misaligned illumination. This is similarly also described by Katz et al. 2009 using a bare fiber approach at various not further specified distances of 0.1 - 2 mm. They report the artifact as monophasic signal which is superimposed with assumed nerve activity and accompanied by both acoustic sounds and visible movement of the nerve. This is in direct contradiction to the biphasic form we encountered and verified by nerve-crush control and thus the effect warrants further independent research and verification. At least two underlying mechanisms come to mind: laser energy conversion by absorption into heating, local thermal nerve shrinking, or a combination of both resulting in an observable movement artifact in the data acquisition system. Heating via water as chromophore is taken as one of the facilitators for the underlying mechanism for INS (Wells et al. 2010, Shapiro et al. 2012). The difference between the here described artifact and other measurements could be explained by differing setups. The nerve bath guarantees an optimal hydration for the nerve, which is preferred to intermittently re-wetting nerve fibers in other ex vivo setups, potentially resulting in an even stronger extent of the effect. As the heating at the end of the laser illumination ceases, the expansion is reverted and could explain the second inverted signal shape. Audible photo-thermal expansion caused indeed visible waves in the nerve bath which might result in a similar signal form, however, we argue that this effect would then be observable in every measurement regardless of successful CAP generation, as it is not reliant on a certain straining precondition of the nerve. Since this is not the case, we discard this mechanism as a cause for the artifact and need to consider it separately from cases where the buffer solution in the nerve bath was irradiated as well.

Another possible mechanism for this movement artifact is local reversible shrinkage along the nerve sheath as has been observed in in vivo studies. Here, the severity appeared to be of increasing magnitude with increased energy deposition (Schlett 2020), which is in line with our observed increasing amplitude of the artifact’s signal at higher radiant exposures. We hypothesize that heat induced contraction in collagen fibres (Wall et al. 1999, Humphrey 2002) of the epineural and perineural layers are the underlying mechanism. The first part of the observed mirrored spikes would be generated through local shrinking and shortening of the whole nerve while the second would occur through strain release. This pull-relaxation cycle could indeed be negated by changing the distance between the two fixating pins in the nutrition bath. It is furthermore not observable when the excised nerve is sufficiently long and the mounting is done without any pre-strain.

It should be noted, that inhomogeneous hydration might exacerbate changes in heat-induced longitudinal strain and thus add another confound to ex vivo INS.

The second prominent artifact was encountered by adjacent or direct illumination of the hook electrode. Similar signals are reported for optogenetics setups where tungsten electrodes were utilised with an illumination at 473 nm (Han et al. 2009). Here, despite using the longer wavelength 1470 nm laser, the erroneous signal is well distinguishable in comparison with a variety of different and longer pulse widths as it scales accordingly in time. Therefore, we utilized multiple control measurements, each with different illumination times. A strong difference to CAPs becomes clearly visible over the course of increasing illumination times.

In lieu of a definitive explanation of the underlying mechanism, a photovoltaic effect with thermo-electric coupling into the tungsten or platinum electrodes will be discussed.

We argue against a true photoelectric effect (Lenard 1902, Einstein 1905) causing the observed artifact signal. Photoelectric effect occurs when electrons are released from a metal (usually in high vacuum) after excitation by photons (Kozai and Vazquez 2016) depending on the receiving material and on the incident photons. The latter need to have sufficiently high energy and thus have to feature short enough wavelengths. For tungsten, the cutoff wavelength amounts to 257 ± 0.5 nm. (Warner 1929), whereas for platinum electrodes it amounts to 196.2 nm (DuBridge 1928). A potential detectable multi-photon photoelectric effect with rate scaling non-linearly with the intensity would require, given our 1470 nm wavelength, an intensity in the range of 10^12^ -10^13^ W/cm^2^ (Supplemental Material 3). This value runs magnitudes above the capabilities of our employed laser, whose intensity is in the range of 10^4^ W/cm^2^. Furthermore, the photoelectric effect occurs in tungsten in the attosecond range (Ossiander et al. 2018). Contrary to that our observed delay is in the millisecond range which does not corroborate a photoelectric effect.

A similar artifact, the photovoltaic (or Bequerel-)effect has been reported in electrodes for the optogenetic field at smaller wavelengths, demonstrating an instantaneous emergence (Kim et al. 2019, Kozai and Vasquez 2016). Unfortunately, our artifact doesn’t appear instantaneously, forcing us to take an additional transient thermal disturbance or a capacitive involvement into account. A thermal disturbance of the metal’s double layer could play a role as a capacitance causing the delay. In any ways, the optically induced electronic signal has to be taken seriously and warrants further investigation in its causes as clearly, care must be taken to ensure that electrodes are not accidentally illuminated by the laser beam. This risk is particularly exacerbated in setups with diverging light beam cones from fibers.

## Conclusion

We report on a new setup for ex vivo infrared nerve stimulation utilizing a continuously perfused nerve bath maintaining reliable nutrient’s supply to an excised rat sciatic nerve. A free, lens-projected beam of a 1470 nm IR-B laser was able to elicit compound action potentials recorded by metal hook-electrodes and measured by an electrophysiology amplifier. Experiments motivated by neuropharmacological exposure studies could not be performed as the nerve’s viability in re-excitability degraded too rapidly at the chosen, physiological temperature. However, we are convinced, that by optimizing both medium and nerve extraction the presented approach may lead to a stable platform enabling to elucidate the still controversially discussed mode of operation of INS.

We furthermore report on prominent artifacts encountered with this INS setup and present ways to identify them to reduce the risk of false positive measurements and to help future researchers to deal with these challenges accordingly.

## Data availability

The experimentally acquired data of this study are available upon reasonable request from the authors.

## Acknowledgments

The research was funded by the Federal Ministry of Research, Technology and Space (BMFTR) in the NeuroPhos Project (grant number 13GW0155C), and DiaQNOS Project (grant number 13N16460).

