## Supplemental Materials for "Ex vivo Infrared Nerve Stimulation on the Rat Sciatic Nerve: Challenges and Pitfalls"

### Supplemental Material

#### 1) Whole pulse train CAPs per pulse width.

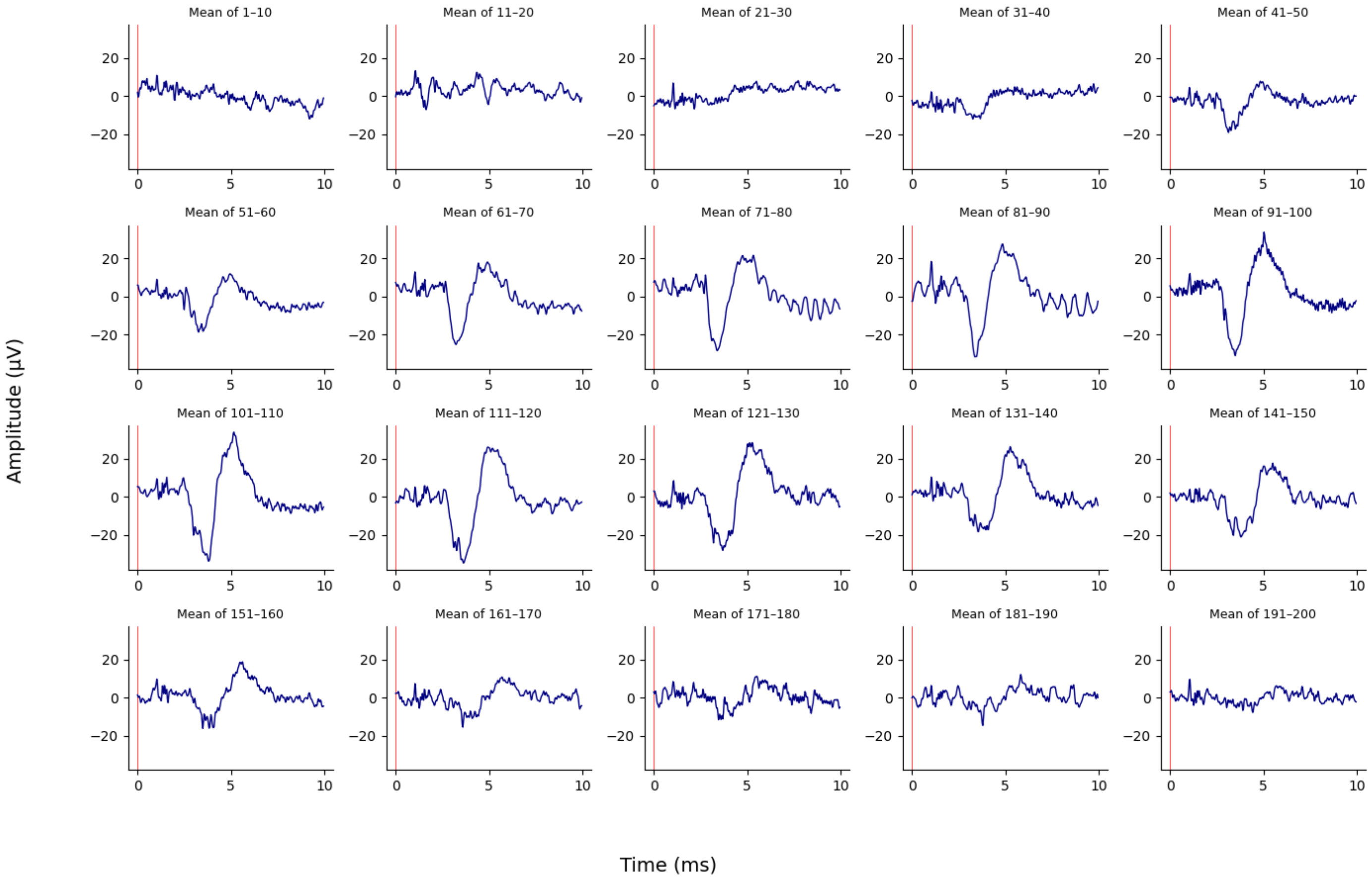


*Fig. 1: Evolution of observed CAP signal means (blue) over 20 pulse sequences. For each 10 single pulses used for averaging one pulse width was used, gradually increasing from 0.1 – 2 ms.*

#### 2) Lens system list

| Lens | Focal length f [mm] | Diameter d [mm] | Amount |
|---|---|---|---|
| Thorlabs LA1116 | 10 | 6 | 1 |
| Thorlabs LA1251 | 100 | 25 | 1 |
| Thorlabs LA1255 | 50.1 | 25 | 2 |

3) Photoelectric effect:

Energy of photons at 1470 nm: $E_\gamma = \frac{hc}{\lambda} = 0.8434\text{eV}$.

Work function for tungsten: $I_P = 4.55\text{eV}$ (Strek et al. 2021)

Therefore, the number of photons needed to overcome the ionization potential is $n = \frac{I_P}{E_\gamma} = \frac{4.55\text{eV}}{0.8434\text{eV}} = 5.394 \approx 6$.

In the strong-field regime, the Keldysh theory becomes the relevant framework to treat photoionization.

Multiphoton ionization (as opposed to tunnel ionization) occurs for a Keldysh parameter (Keldysh 1965) $\gamma \gg 1$, where: $\gamma = \sqrt{\frac{I_p}{2U_P}} = \frac{\omega\sqrt{2m_e I_p}}{eE}$, but sets on already for $\gamma \gtrsim 1$ in the non-adiabatic perturbative regime.

For the laser with a radiant exposure of 25.71 J/cm² (highest used in the presented experiments) at 2 ms pulse width, and a focal point diameter of d = 352 µm the peak power is calculated as $P = \frac{E}{t} = \frac{0.02474J}{0.002s} = 12.37W$. With this the intensity per pulse can be calculated as $I = \frac{P}{A} = \frac{12.37W}{9.73\cdot 10^{-4}cm^2} = 1.27 \cdot 10^4 \frac{W}{cm^2}$. The Keldysh parameter we obtain for these settings at 1470 nm is on the order of $10^5$ which is well in the perturbative regime. The ionization rate, however, scales non-linearly with the intensity $W^{(n)} = \sigma^{(n)} I^n$ and for typical n-photon cross-sections the rate is negligibly small. Compared to the highest intensities, still in the multiphoton regime ($\gamma \gtrsim 1$), on the order of $10^{13} \frac{W}{cm^2}$, the ionization rate at the highest radiant exposure with $I \propto 10^4 \frac{W}{cm^2}$ would be $10^{54}$ orders of magnitude smaller and therefore practically not observable.
